# A COMPARATIVE STUDY OF ANAESTHETIC AGENTS ON HIGH VOLTAGE ACTIVATED CALCIUM CHANNEL CURRENTS IN IDENTIFIED MOLLUSCAN NEURONS

**DOI:** 10.1101/2020.12.17.423182

**Authors:** Terrence J. Morris, Philip M. Hopkins, William Winlow

## Abstract

1. Using the two electrode voltage clamp configuration, a high voltage activated whole-cell Ca^2+^ channel current (I_Ba_) was recorded from a cluster of neurosecretory ‘Light Yellow’ Cells (LYC) in the right parietal ganglion of the pond snail *Lymnaea stagnalis*.
2. Recordings of I_Ba_ from LYCs show a reversible concentration-dependent depression of current amplitude in the presence of the volatile anaesthetics halothane, isoflurane and sevoflurane, or the non-volatile anaesthetic pentobarbitone at clinical concentrations.
3. In the presence of the anaesthetics investigated, I_Ba_ measured at the end of the depolarizing test pulse showed proportionally greater depression than that at measured peak amplitude, as well as significant decrease in the rate of activation or increase in inactivation or both.
4. Within the range of concentrations used, the concentration-response plots for all the anaesthetics investigated correlate strongly to straight line functions, with linear regression R^2^ values > 0.99 in all instances.
5. For volatile anaesthetics, the dose-response regression slopes for I_Ba_ increase in magnitude, in order of gradient: sevoflurane, isoflurane and halothane, a sequence which reflects their order of clinical potency in terms of MAC value.

## INTRODUCTION

General anaesthetics form a chemically diverse group of agents which have in common the property of inducing narcosis and analgesia to varying degrees. They tend to depress neuronal excitability and synaptic transmission, though detailed responses vary for the particular drug and are often cell specific. Work with both vertebrate and invertebrate neuronal preparations has shown both [Ca^2+^]_i_ (intracellular calcium concentration) and Ca^2+^ influx to be dramatically affected by a number of anaesthetics. Increases in [Ca^2+^]_i_ in the presence of volatile anaesthetics has been shown in CA1 hippocampal cells in rats (Mody *et al*., 1991) and cultured molluscan neurons in the absence of extracellular Ca^2+^ (Ahmed et al, 2020; Winlow *et al*., 1995), implying an anaesthetic triggered release of Ca^2+^ from intracellular stores. Voltage-gated calcium channels within the cell membranes also appear to be important targets for these agents and calcium influx has been shown to decrease in a dose-dependent manner in their presence (Yar & Winlow, 2016). T-type calcium currents (see below) have also been shown to be differentially sensitive to volatile anaesthetics at clinical concentrations in various cell types (McDowell *et al*., 1999). These are interesting findings but the dual action of Ca^2+^ adds complexity to understanding global effects on cell activity. Ca^2+^ is also a major second messenger and plays a central role in the control of a variety of cell processes, so that alterations of calcium influx and [Ca^2+^]_i_ have the potential to modify numerous, interrelated neuronal functions. Transient increases in [Ca^2+^]_i_ may be initiated by release from intracellular storage sites associated with the endoplasmic reticulum (Yamaguchi, 2019) and/or an increase in Ca^2+^ permeability of the plasma membrane associated with neural activity or the actions of neurotransmitters (Miller, 1991) and may operate by a calcium-induced calcium release mechanism (Sandler and Barbara, 1999; Petrou et al, 2017).

In vertebrate tissues, calcium channels have been classified by their electrophysiological and biochemical properties. Nowycky *et al*. (1985) identified three types of Ca^2+^ channel in whole-cell currents recorded from chick dorsal root ganglion cells, called L-type, T-type and N-type. They have become the prototypes that form the basis of a widely accepted system of calcium channel classification (Catterall, 2011; Catterall, et al, 2019, 2020). The transient T-type current is activated by small depolarisations, termed Low-Voltage-Activated (LVA), whilst sustained L-type currents are activated by large depolarisations - High-Voltage-Activated (HVA) - and blocked by organic calcium channel antagonists. N-type currents are also included in the HVA category but can be transient or sustained. An increasing number of calcium channels from both vertebrate (Hoehn *et al*., 1993) and non-vertebrate systems (Pearson *et al*., 1993) have proven difficult to identify within this scheme. Molluscan neurons in particular can display HVA Ca^2+^ currents with conflicting pharmacological and electrophysiological profiles (Kits & Mansvelder, 1996), but L-type currents have clearly been identified as the sole HVA current in isolated, cultured pedal I cluster neurons of *Lymnaea stagnalis* (Yar and Winlow, 2016).

The molluscan CNS (central nervous system) has several specific advantages for the neurobiologist exemplified in *Lymnaea* (Kerkut, 1989; Leake & Walker, 1980); nerve cells are accessible, relatively simply organised, easily identified and in many cases large enough for two electrode work. Moreover, volatile anaesthetics produce many changes in behavior and neuronal activity in *Lymnaea* that equate well to those found in mammals (McCrohan et al, 1987; Winlow & Girdlestone, 1988; Girdlestone et al 1989) and facilitate its use as a single “model” system in studies on anaesthetic mechanisms at behavioral and cellular levels (Winlow, 1984; Winlow, et al, 2018; Moghadam et al, 2019).

The HVA calcium currents recorded from the cultured pedal I cluster neurons of *Lymnaea* (Yar and Winlow, 2016) using single electrode voltage clamp showed a reversible, dose-dependent suppression of Ba^2+^ mediated current by halothane at concentrations ranging between 0.5 to 4.0 percent. Here we report on data using the two-electrode voltage clamp technique to investigate calcium channel currents recorded from the Light Yellow Cell (LYC) group of neurosecretory neurons in the right parietal ganglion of *Lymnaea* in the intact brain, which play a general role in body fluid regulation (Benjamin and Kemenes, 2020). They lie on the ventral lobe of the right parietal ganglion and can be observed from either the dorsal or ventral surface of the ganglion. LYCs have large somata and fire spontaneous bursts of spikes (van Swigchem, 1979). Their action potentials have a prominent shoulder or pseudoplateau phase, believed to be largely Ca^2+^ driven (Aldrich, Jr. *et al*., 1979; van Swigchem, 1979).

Studies of the effects of general anaesthetics on voltage gated Ca^2+^ channels have often involved the use of a variety of preparations and cell type, frequently at concentrations outside the clinical range. The purpose of this study was, therefore, to determine the action of a number of general anaesthetics, at clinical concentrations, on calcium currents recorded from the same identified cell group in a single model system. In this way the dose-response profile and relative potency of each agent can be directly compared. Here, we characterize the electrophysiology and pharmacology of the LYC Ca^2+^ currents, and the effects upon them of clinical concentrations of the volatile anaesthetics halothane, isoflurane and sevoflurane and the systemic anaesthetic sodium pentobarbital, which is now mostly used in veterinary anesthesia (Lester et al, 2012).

## METHODS

Snail brains were prepared according to the methods of Benjamin & Winlow (1981) in a HEPES buffered snail saline (see below) at room temperature, approximately 20° C. Briefly, intact central ganglia were pinned out in the perfusion chamber, ventral surface uppermost, and the sheath of connective tissue removed from the right parietal ganglion with fine ground forceps. The inner cell integument was softened with several drops of protease solution (Pronase from *Streptomyces griseus*, Boehringer Mannheim Biochemica, in a solution of 4 mg/ml of snail saline) applied directly to the ganglia, and washed off thoroughly with saline after about 3 minutes. The preparation was then perfused continuously at 3.5 ml per minute with aerated saline. After 10-15 minutes the perfusing medium was switched to the recording solution, also at room temperature, after which a cell within the LYC cluster was selected for microelectrode penetration.

### Solutions

Preliminary dissection and perfusion prior to recording was carried out in a snail saline (Benjamin & Winlow, 1981) containing (mmol l^-1^): Na^+^ 59.4, K^+^ 2.0, Mg^2+^ 2.0, Ca^2+^ 4.0, Cl^-^ 38.0, HEPES (4-(2-hydroxyethyl)-1-piperazineethanesulfonic acid) 50.0, and glucose 0.3. The pH was corrected to 7.8 with 2M NaOH.

All current measurements were made from cells in the intact brain, bathed in recording solution which contained (mmol l^-1^): Na^+^ 6.5, Cs^+^ 2.0, Mg^2+^ 1.5, Ba^2+^ 10, Cl^-^ 23, Br^-^ 30, tetraethylammonium (TEA) 30, 4-aminopyridine 10, HEPES 10, glucose 5 and pyruvate 10. The whole cell currents measured were Ba^2+^ currents rather than Ca^2+^ currents *per se*. Ba^2+^ was used as the charge carrying ion since no currents were detectable when Ba^2+^ was replaced by Ca^2+^. Possible explanations for this are first, that the calcium channels in these cells are much more permeant to Ba^2+^ than to Ca^2+^, thereby increasing the signal to noise ratio to a resolvable level and, second, Ca^2+^ entering the cell might produce Ca^2+^-dependent inactivation of the channels being studied. Ba^2+^ also has the advantage that it blocks contaminating K^+^ currents, which were further blocked by the presence of TEA^+^ and Cs^+^ in the recording solution. [Na^+^]_out_ was kept low to exclude voltage-dependent Na^+^ currents. The addition of pyruvate was found to increase the recording time of cells considerably. Organic Camchannel blockers, verapamil (Sigma) and nifedipine (Sigma), were dissolved initially in ethanol to give stock solutions of 1 mg ml^-1^ before being diluted in the recording solution to the required concentration.

The pipette filling solution, modified from Orchard *et al*. (1991), contained (mmol l^-1^): KCl 2500, calcium buffer ethyleneglycol-bis (-amino-ethyl ether) N,N’-tetra-acetic acid (EGTA) and K-ATP 10.

### Administration of anaesthetics

Volatile anaesthetics were vaporized into air using Ohmeda vaporizers set to the desired percentage mixture. The anaesthetic-air mixture was bubbled at 1 litre per minute into a small volume (about 150 ml) of recording solution in a 250 ml flask and vented though a closed extraction system. Percentage vapour concentration was routinely calibrated using a Normac anaesthetic agent monitor. Equilibration time was taken as 15-20 minutes (Girdlestone et al., 1989) after which a stopper containing an underwater seal air inlet was inserted and the solution used immediately via a closed delivery system. Millimolar concentrations of isoflurane at 1, 2 and 4% were determined by gas-liquid chromatography as (mean ± SD for n = 3): 0.44 ± 0.05, 0.76 ± 0.18 and 1.89 ± 0.37 respectively (in an ideally equilibrated solution a 1% vol/vol concentration of gas represents 0.42 mM). These values approximate those for equilibrated halothane solutions (Winlow et al., 1998) and are very close to the normal clinical concentration of halothane in arterial blood (Davies et al., 1972).

The clinical use of pentobarbitone in humans is nowadays rare, although it may be used in veterinary anesthesia (Lester et al, 2012). Therefore, most uptake studies are on barbiturates used in contemporary anaesthesia, thiopentone sodium in particular. Thiopentone sodium has the same clinical potency as pentobarbitone (Dundee, 1974) but possesses contrasting and more desirable pharmacokinetic properties (Lant, 1982). The concentration range of pentobarbitone used in this investigation was 200 to 800 μM, which equates to approximately 50 to 200 μg per ml of recording solution. The median dose of thiopentone required to induce anaesthesia is about 3.5 mg/kg body weight, equivalent to approximately 100μg/ml plasma concentration (Dundee *et al*., 1982).

### Recording from cells

Micropipettes (resistance 8–10 MΩ) were pulled on a one stage vertical puller using filamented borosilicate glass (Clarke Electromedical Instruments). An Axoclamp 2A amplifier was used to clamp cells and acquire data via proprietary software (PClamp 6.0) running on a 486 DX 33MHz IBM clone PC through an Axoclamp Digidata 1200 digital-analogue interface. Data were sampled at 5 kHz, with automatic leak subtraction, estimated electronically before each test pulse as the sum of currents developed during a series of four inverted pulse steps at ¼ amplitude of the test pulse itself. A Gaussian filter with 500 Hz cut-off was applied to the raw data after acquisition to attenuate noise.

Optimum results were achieved when the preparation was allowed to equilibrate in recording solution 4–6 minutes before inserting microelectrodes into the selected cell. Passive electrical parameters of the cell membrane were measured by manually injecting current. Mean values for initial τ and input resistance were, respectively (mean ± S.E. for n = 25): 78.3 ± 5.6 ms, 20.7 ± 1.7 MΩ. Useable cells had an initial input resistance >10 MΩ and τ of approximately 50 ms, although these values could increase by up to a factor of 3 once the cell was clamped to the holding potential of −70 mV. At the holding potential, calcium channel current was monitored in response to single depolarizing pulses to 0 mV, and would gradually increase in magnitude over several minutes, reaching a stable value of between 40 to 150 nA, depending on cell size. From the same holding potential, current–voltage (I–V) curves were constructed by applying depolarizing voltage pulses (duration 70 or 150 ms) over a range of test potentials in increments of 7 or 10 mV.

For experiments involving the application of drugs I–V pulse protocols were first performed in the control situation at a holding potential of −70 mV. During perfusion with the control solution and subsequently the test solution, a series of unitary depolarisations to 0 mV were applied. When the calcium channel current re-stabilised another I–V curve was obtained as above. Between applications of the same drug at different concentrations, the preparation was washed with control solution until the calcium current amplitude closely approximated the control value.

### Statistical analysis

Data are expressed as mean ± standard error as appropriate

## RESULTS

### 1) The properties of whole cell calcium channel currents recorded from LYCs

Two electrode voltage clamp recordings from LYCs in *Lymnaea* revealed large inward currents elicited by depolarizing the cells from a holding potential of −70 mV in the recording solution with Ba^2+^ as the charge carrier. Figure 1A shows a selection of current traces recorded from a large LYC. The inward current (I_Ba_) was elicited during stepped depolarisations of 70 ms, to test potentials between − 50 and + 48 mV in 7mV steps. Currents disappeared when Ba^2+^ was omitted from the external solution, but increased in amplitude as [Ba^2+^]_out_ increased (not shown). I_Ba_ was neither affected by nifedipine at 50 μM nor verapamil at 20 μM but was completely abolished by Cd^2+^ at 100μM. At small depolarisations the current is slow to activate, but the activation rate increases in step with larger depolarisations. Inactivation is slow, and partial, but increases slightly in rate with larger depolarisations. These observations are summarized in Table 1, which gives the time constant for activation and the time to peak for weak and strong depolarisations, both of which are larger for test pulses to −30 mV than to 0mV.

**Figure 1.**
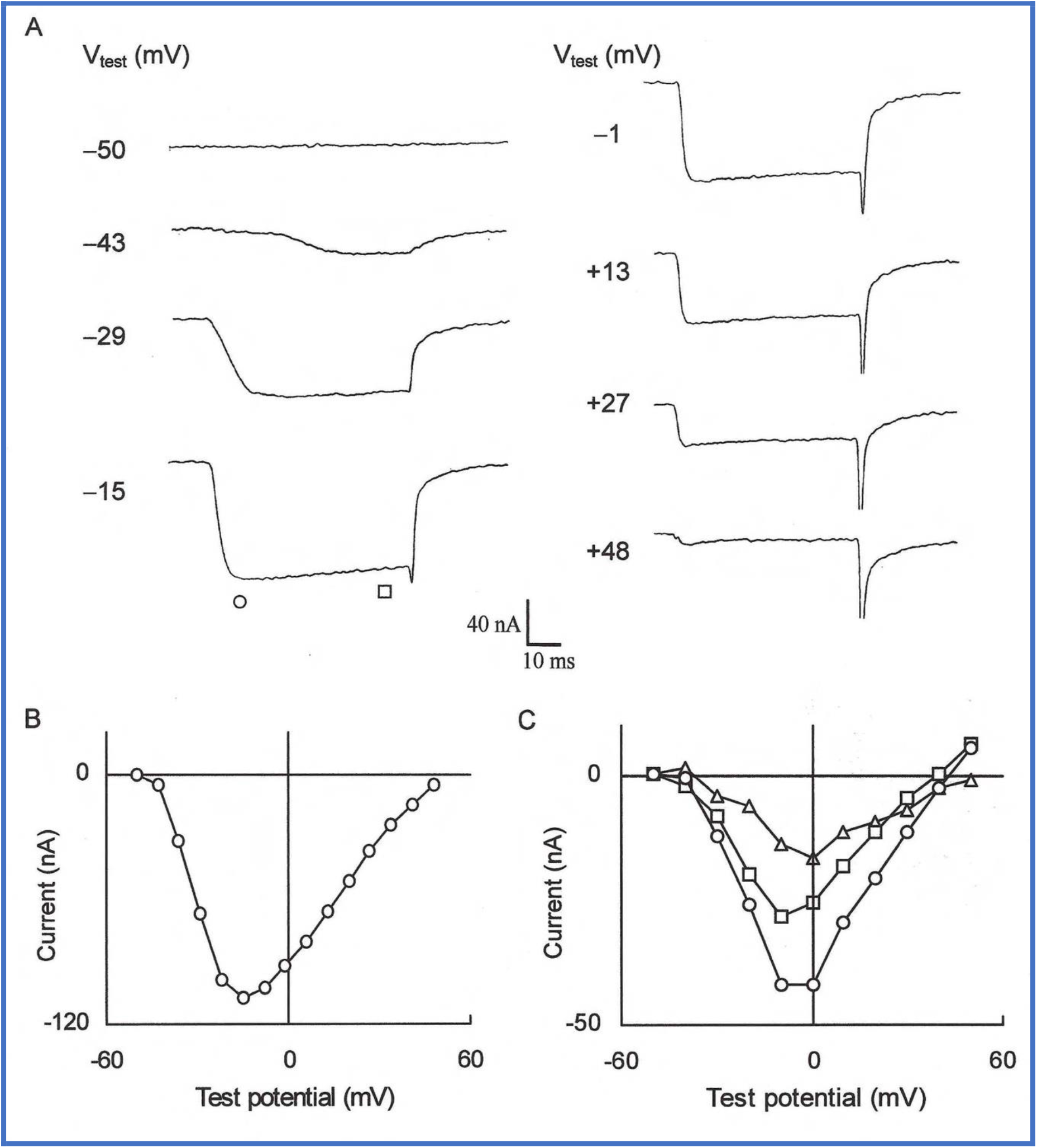
A) Whole cell Ba^2+^ currents recorded from a LYC neuron in *Lymnaea stagnalis* held at −70mV and step depolarised for 70 ms to the test potentials indicated next to each current trace. (B) I–V curve of peak amplitude of currents in ‘A’. (C) I–V curves for peak (○), sustained (□) and transient (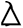 current measurements recorded from a different LYC, step depolarised for 150 ms at each test potential in 10 mV increments.

**Table 1.** Activation and inactivation parameters of I_Ba_ recorded from LYC in response to stepped depolarisations of 150 ms from the holding potential of −70mV.

| Test Pulse (mV) | Time to peak (ms ± S.E.) [n] | Time constant ($\tau$) for activation (ms ± S.E.) [n] | Time constant ($\tau$) for inactivation (ms ± S.E.) [n] |
| --- | --- | --- | --- |
| -30 | 39.6 ± 3.40 [6] | 9.18 ± 3.8 [4] | 92.4 ± 8.5 [4] |
| 0 | 19.90 ± 1.06 [6] | 3.85 ± 0.307 [5] | 44.78 ± 5.63 [4] |

In Figure 1B, the amplitude measured at the peak of each trace is plotted against test potential, to show the current voltage (I–V) relationship for peak current. Inward current first appears at about −45 mV and increases in amplitude with stronger depolarisations to a maximum (I_Ba(max)_) at −15mV, and appears to reverse at about +50 mV. Considering the bimodal activation kinetics observed in current traces, the co-existence of discrete, multiple current components with different activation kinetics could not be excluded. Therefore, in another LYC, I_Ba_ was measured at the peak (I_peak_) and near the end of 150 ms test pulses, when current amplitude becomes sustained (I_sus_). Shown in Figure 1C are I–V plots for I_peak_ and I_sus_ and the measured difference between them (I_trans_), effectively the amount of current that inactivates during a complete step. To characterize these components further, steady state inactivation was explored by varying the holding potential prior to depolarizing the cell to approximately I_Ba(max)_. The plots of I_Ba(max)_ for I_trans_ and I_sus_ against holding potential fit quite well to a single sigmoidal shaped curve (Figure 2) and there were no significant differences between the level of steady state inactivation of any of these currents at each holding potential. Both sustained and transient components begin to activate at −70mV, are 50% inactivated at a holding potential of 41 mV and are fully inactivated when the holding potential reaches −20 mV. Therefore, variation of the holding potential does not separate transient and sustained components of I_Ba_.

**Figure 2.**
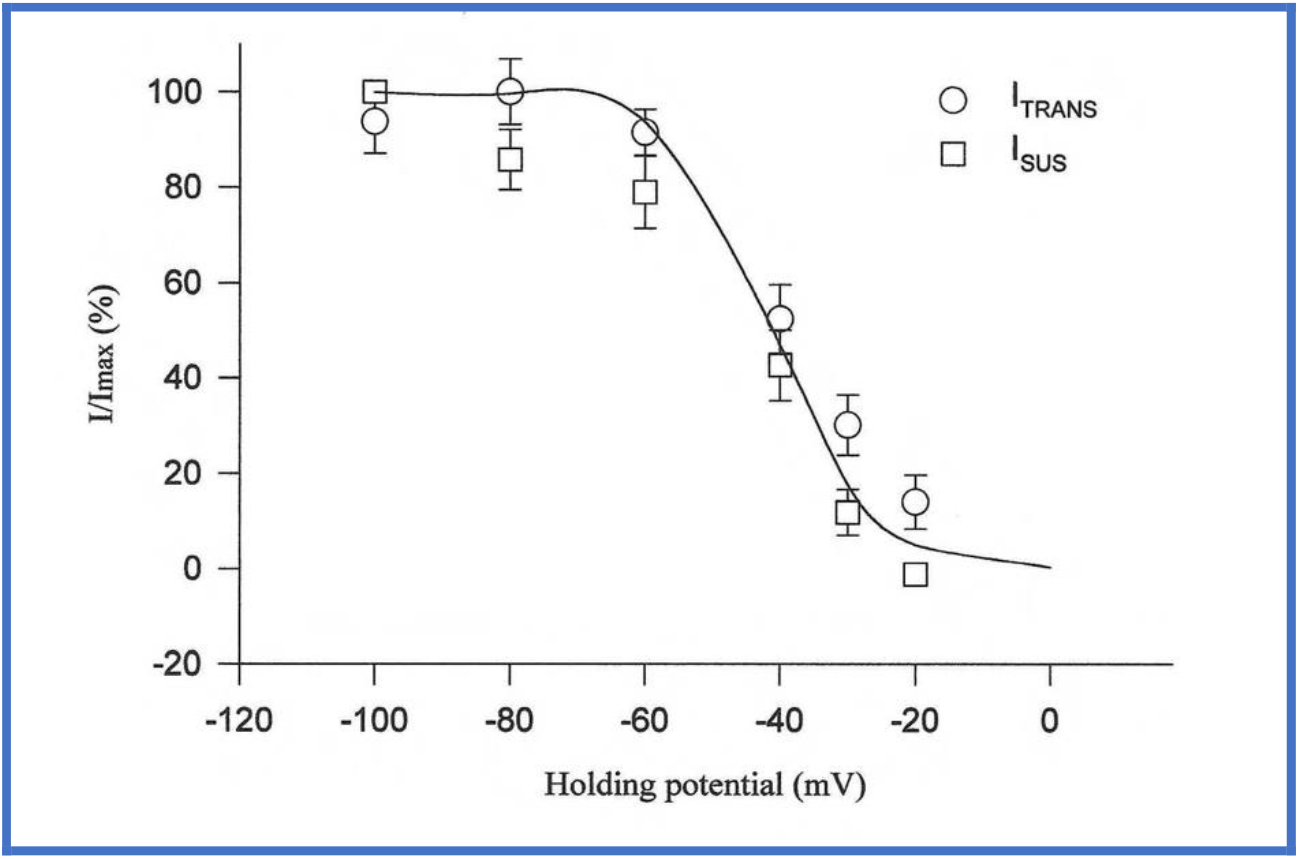
Steady state inactivation curve for I_sus_ and I_trans_, recorded from 8 cells. Currents are expressed as a percentage of the corresponding transient and sustained currents at the holding potential of −100mV for a test potential of −15mV. Holding potentials were (mV) −100, −80, −60, −40, −30, and −20. No significant differences were found between transient and sustained values at each holding potential, using a paired t-test. A single curve was fitted to all data points by the least squares method using the Boltzmann distribution: I/Imax = 1/[1+exp(Vh–V1/2)/k] where V1/2 = −40.88mV, Vh is the holding potential and k = 7o08 mV.

### 2) The effect of anaesthetics on I_Ba_

Calcium channel currents recorded from LYCs were reversibly depressed in the presence of all the anaesthetics tested. For example, Figure 3A shows the effect of halothane on current traces of I_Ba(max)_ recorded from a single LYC. When halothane is applied at 1, 2, and 4%, current amplitude is reduced for the duration of the depolarizing pulse in a dose dependent manner. Figure 3B is a time course plot of peak and end-pulse current elicited by unitary depolarisations to 0 mV recorded during a typical halothane experiment, showing these effects to be fully reversible. In all cases the current at peak amplitude was less sensitive to anaesthetic than that recorded at the end of the depolarizing pulse (compare dose-response plots in figure 4B). There was also a tendency for activation rates to decrease and inactivation to increase instep with increasing anaesthetic concentration. For example, at the highest concentrations used, activation rates were significantly decreased for halothane, isoflurane and pentobarbitone and inactivation significantly decreased for isoflurane sevoflurane and pentobarbitone (Table 2). Therefore, in addition to a reduction in whole-cell calcium channel current, it is clear that these drugs have differential effects on calcium channel gating kinetics.

**Figure 3.**
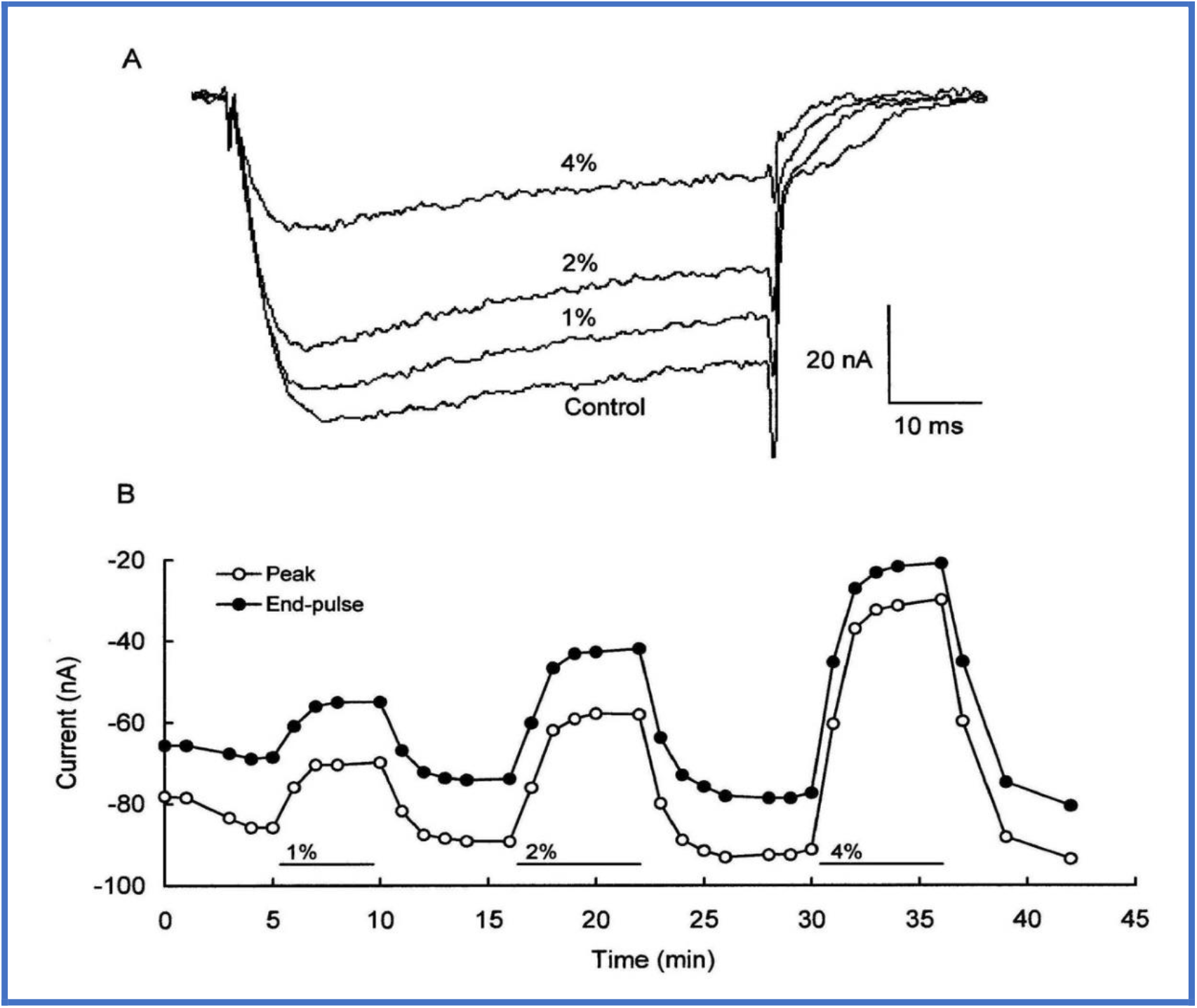
A) Dose-dependent reduction of I_Ba_ by halothane. Halothane was applied successively at 1, 2 and 4% to a single LYC neuron with washes in between. Peak current was monitored and standard I–V pulse protocols were run during the plateau of maximal drug response. Superimposed currents represent I_Ba(max)_, measured as −8mV in this instance for both the control and test conditions. B) Time course of peak and end-pulse I_Ba_ currents recorded from an LYC during a typical experiment in response to unitary 70 ms depolarising pulses to −0mV from a holding potential of −70mV. Halothane was introduced at the concentration and duration indicated by horizontal bars.

**Figure 4.**
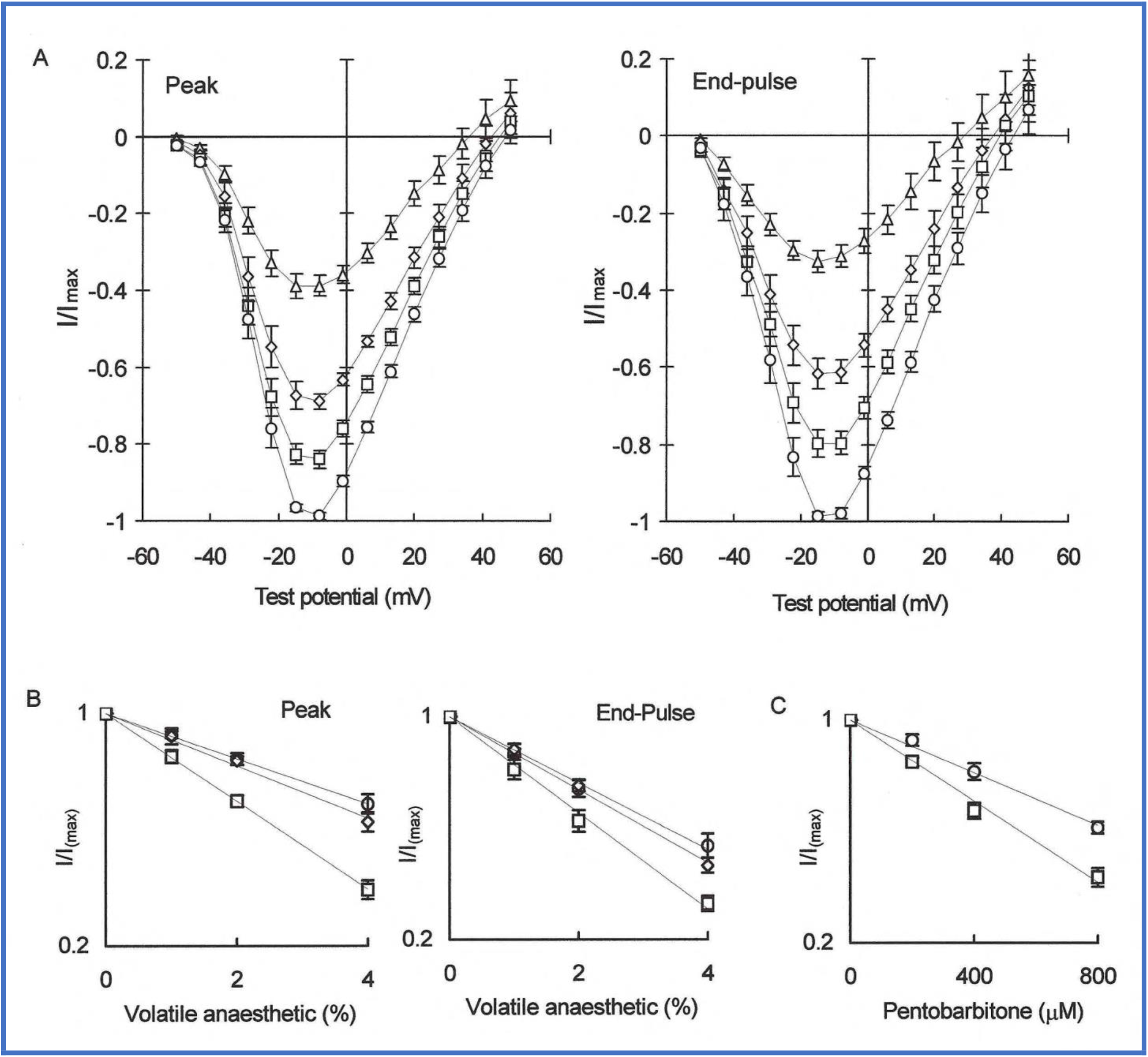
A) The effect of halothane concentration on I–V curves of mean peak and end-pulse I_Ba_ normalised to control I_Ba(max)_ and showing ± S.E.M. as vertical bars. For each trial halothane was introduced successively at 1% (□), 2% (◊) and 4% (∆), interspersed by washes of 2-3 minutes which reversed the effects completely (control ○). I_Ba(max)_ was monitored throughout and pulse I–V protocols performed during the plateau of maximal response to the drug. B) Normalised means of I_Ba(max)_ for peak and end-pulse currents are plotted against percentage volatile anaesthetic concentration for halothane (□), isoflurane (◊) and sevoflurane (○). Vertical bars indicate s.e.m. where n = 6 for halothane and n = 7 for isoflurane and sevoflurane. C) Normalised means of I_Ba(max)_ for peak (○) and end-pulse (□) currents plotted against pentobarbitone concentration. Linear regressions are fitted though each dose-response plot in 4B and 4C using the least squares method. All have an R^2^ value greater than 0.99.

**Table 2.** Mean maximum slope of I_Ba_ traces, measured before (activation) and after (inactivation) the slope turning point at peak amplitude. Volatile anaesthetics were applied at 4% and pentobarbitone at 800 μM. Traces were smoothed with a Gaussian filter at 300 Hz cut-off before taking measurements. Significant statistical differences of slopes compared to control values (marked with an asterisk) were evaluated using Student’s two tailed, paired t-test at the 5% level, n = 5 for each data set. Differences in control values reflect variations in average animal size, and hence LYC size, of the batch of samples.

| | Mean activation slope (nA/mV) ± S.E | | Mean inactivation slope (nA/mV) ± S.E | |
| --- | --- | --- | --- | --- |
|  | Control | Drug | Control | Drug |
| Halothane | $-12.87 \pm 2.44$ | *$-6.45 \pm 1.34$ | $0.525 \pm 0.070$ | $0.688 \pm 0.085$ |
| Isoflurane | $-13.31 \pm 1.38$ | *$-9.19 \pm 1.34$ | $0.529 \pm 0.079$ | *$0.848 \pm 0.065$ |
| Sevoflurane | $-6.93 \pm 0.99$ | $-6.00 \pm 1.35$ | $0.288 \pm 0.038$ | *$0.464 \pm 0.085$ |
| Pentobarbitone | $-10.13 \pm 1.58$ | *$-6.95 \pm 1.19$ | $0.515 \pm 0.073$ | *$0.722 \pm 0.148$ |

### 3) The effect of anaesthetics on the current-voltage relationship of I_Ba_

To determine the effect of drugs on the current-voltage relationship, I–V plots were constructed for mean peak and end-pulse current recorded from several cells and normalized to control values of I_Ba(max)_. Thus, for halothane, two families of I–V curves were produced, each for control conditions and for halothane at 1, 2 and 4% (Figure 4A). These show that increasing halothane concentration reduces I_Ba_ in a dose-dependent manner across the entire voltage range, with end-pulse current being slightly more sensitive to the anaesthetic. The I–V curve shape is well preserved across the voltage range except at the most positive test potentials when anaesthetic concentration was high. Here, net outward current increases, especially for end-pulse current where Erev has fallen from 40 to 25 mV. This effect was less apparent for other anaesthetics tested but could indicate either that calcium channel kinetics are altered in direct response to the anaesthetic or that an unspecified repolarizing current, such as a K^+^ channel current, was being activated at the higher drug concentration.

Similar families of I–V curves were produced for isoflurane, sevoflurane and pentobarbitone, which differed principally in their dose-response profile. These are summarized as dose-response plots of I_Ba(max)_ in Figure 4B for volatile anaesthetics and 4C for pentobarbitone (R^2^ > 0.99 in all instances). Therefore, I_Ba_ is reduced in direct proportion to the concentration of anaesthetic for both the volatile and non-volatile drugs within the concentration range of drugs used. The proportional decrease in I_Ba(max)_ for a given concentration of drug is greater for end-pulse currents which have steeper dose-response slopes. For the volatile anaesthetics, halothane produces the greatest depression of I_Ba_, followed by isoflurane, and then sevoflurane. The slope coefficients for peak currents show wider variation compared to those of end-pulse currents, where the regression slopes are divergent. Hence, the relative ability of the volatile anaesthetics to depress I_Ba_ are different when comparing peak and end-pulse I_Ba(max)_ regressions of dose-response. Sensitivity of I_Ba_ to pentobarbitone in the range 0.2 - 0.8mM is about twice that for volatile agents, which had an equivalent dose range of approximately 0.4 – 1.6 mM. Pentobarbitone, a compound physically and chemically distinct from volatile anaesthetics, also has a linear dose-response profile at these concentrations.

The anaesthetics investigated produced a reversible dose-dependent depression of I_Ba_ in LYCs of *Lymnaea stagnalis* and, in all cases, current at peak amplitude was less sensitive than that recorded at the end of the depolarizing pulse. Most of the drugs tested produced a significant decrease in activation rate and an increased inactivation rate. Plots of I_Ba(max)_ against anaesthetic concentration correlate highly to linear functions so that the potency of the drug to depress I_Ba_ is proportional to the straight-line slope of the regression.

## DISCUSSION

The aim of these experiments was to compare the effects of a number of general anaesthetic agents on whole cell calcium channel currents recorded from identified light yellow cells, located in the central ganglia of *Lymnaea stagnalis*. The results show that these agents alter calcium channel function, represented as the dose-dependent depression of a calcium channel current, often associated with a decreased rate of activation and accelerated inactivation. Moreover, use of a single model system provides a reliable assay for comparing the relative effects of drugs. In solutions containing Ba^2+^ as the charge carrying ion, large inward currents were observed when cells were depolarized. That these currents are carried though Ca^2+^ channels is supported by the following: (1) The experiments were performed under conditions designed to eliminate as far as possible currents carried by ions other than Ca^2+^ or Ba^2+^ (see Methods). (2) Any increase or decrease in [Ba^2+^]_out_ produced a corresponding change in I_Ba(max)_, whilst excluding Ba^2+^ ions from the recording solution reversibly obliterated I_Ba_. (3) The current was blocked by the calcium channel blocker Cd^2+^ though not by nifedipine or verapamil. I_Ba_ appears to have two HVA components, I_sus_ and I_trans_, which were indistinguishable by their steady state inactivation kinetics and the pharmacological probes used in this study. Molluscan neuron calcium channel identification cannot always be made on the basis of the L-, N-, and T-type currents seen in vertebrate neurons. The resistance of I_Ba_ to the dihydropyridine nifedipine makes it unlikely that any L-type channel currents were present in these cells, although they were previously demonstrated in cultured pedal I cluster neurons of *Lymnaea* (Yar and Winlow, 2016). In addition, the time constant for I_Ba_ inactivation (τ ≈ 50-90 ms) is relatively fast for L-types, more within the range of N-type calcium channels, to which I_Ba_ is otherwise dissimilar. Compared to N-types I_Ba_ possesses an activation range too negative, and a holding potential too positive for their complete steady state inactivation. Pharmacological characterization of I_Ba_ using more specific calcium channel blockers, such as the *ω*-conotoxins and *ω*-agatoxins, would therefore be of interest.

Halothane produced the steepest dose-response slope, followed by isoflurane and then sevoflurane. The order of potency of these anaesthetics to depress I_Ba_ reflects their clinical order of potency, in terms of MAC value (the minimum alveolar concentration required to induce anaesthesia in 50% of the population)(Nickalls and Mapleson, 2003), illustrated in Figure 5. This is a remarkable finding, perhaps signifying the importance of HVA Ca^2+^ channel responses in anaesthetic action, as well as endorsing the suitability of *Lymnaea* CNS as a model system in studies of this type. However, there are complex tissue and channel specific variations of response to general anaesthetics, which appear to exclude a simple unified mechanism of action. In this respect these results contrast sharply with those of McDowell *et al*. (1999) for LVA T-type currents in dorsal root ganglion neurons and two neuroendocrine cells, with isoflurane and enflurane being more potent than halothane.

**Figure 5.**
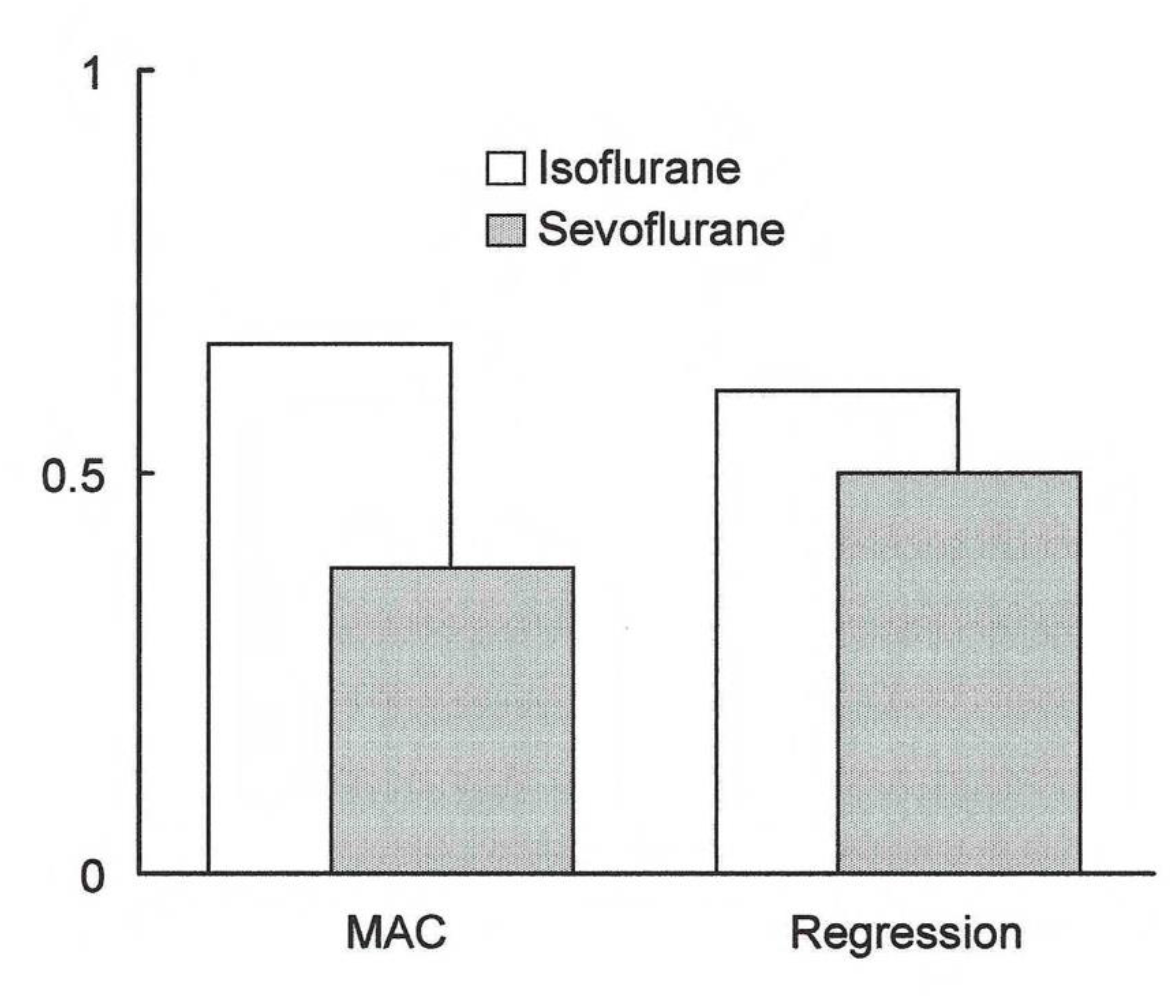
The relative potency of Isoflurane and Sevoflurane in proportion to halothane: to suppress I_Ba_ (as the gradient of dose-response regression slopes) (at right) and when expressed as MAC (at left). Values are normalised to halothane, which is taken as unity.

Pentobarbitone also produces dose-dependent depression of I_Ba_ recorded from LYCs, even though barbiturates are physically and chemically distinct from volatile agents. Similar depression of the HVA human recombinant α1a P/Q type channels by pentobarbital has been demonstrated elsewhere (Schober et al 2010) and involves slow open channel block. However, both sets of findings would appear to cast doubt on some physiochemical theories of anaesthetic action, for example, the correlation between gas-lipid solubility and anaesthetic potency (For a discussion of physiochemical theories see Halsey,1989), at least in terms of effects on voltage gated calcium channels. However, the discovery of soliton pulses associated with action potentials (Tasaki and Iwasa, 1982; Heimberg and Jackson, 2005; Johnson and Winlow, 2018) may yet provide an alternative explanation of anaesthetic actions (Winlow and Johnson, 2018) and may yet unify the modern lipid and membrane protein hypotheses of anaesthesia.

The fact that the HVA currents we have reported here are insensitive to both nifedipine and verapamil strongly suggest that they are LCa_v_2 type channels originally described in *Lymnaea* VD4 neurons. (Haung et al, 2010), but further work will be required to demonstrate this with certainty. Unlike mammalian Ca_v_2.1 and Ca_v_2.2 channels, the *Lymnaea* channels are inhibited slowly by G-protein dependent activation of cAMP, whereas activity-dependent G-protein regulation has evolved in vertebrates, where the G-proteins contain βγ subunits which are missing in *Lymnaea*.

The suppression of voltage-dependent calcium currents is likely to have multiple effects on neuronal function. For example, P/Q type currents are thought to be carried via Ca_v_2.1 channels and are directly involved with synaptic vesicle exocytosis, which is inhibited by isoflurane in rat dopaminergic midbrain neurons, but non-dopaminergic neurons were insensitive to isoflurane (Toturo et al, 2019). However, barbiturates are also known to inhibit noradrenaline and dopamine release with the P/Q voltage sensitive calcium channels (Hirota et al, 2000). Moreover, many secondary messenger pathways, including Ca^2+^ channel activity itself, are modulated by [Ca^2+^]_i_. The ability of ATP to arrest HVA Ca^2+^ channel current rundown, observed here and elsewhere, supports the idea of phosphorylation being central to their maintaining their activity (Chad *et al*., 1987). Elements of protein kinase mediated regulatory pathways are feasible targets for general anaesthetics given the heterogeneous effects these agents have on cells. Firestone & Firestone (1989) found that clinically relevant concentrations of various structurally heterogeneous general anaesthetics reduced PKC binding to an activating ligand in a concentration-dependent manner.

Voltage-dependent Ca^2+^ influx supports Na^+^ mediated depolarisations, so one would expect cell excitability to be reduced by the Ca^2+^ blocking effects of the agents. It has been shown, using the same model system, that a calcium-dependent pseudoplateau of action potentials is abbreviated in the presence of halothane (Winlow *et al*., 1982; Winlow *et al*., 1989; Winlow *et al*., 1992). Thus a briefer and impoverished Ca^2+^ influx is represented at synapses and, since calcium influx is a crucial event in neurotransmitter release (Augustine *et al*., 1987), synaptic transmission is likely to be inhibited. Investigations on *in vivo* and reconstructed synapses between *Lymnaea* neurons have revealed that halothane affects both inhibitory and excitatory responses of pedal A cluster neurons challenged by FMRFamide or electrical stimulation of the pre-synaptic cell (Spencer *et al*., 1995; Spencer *et al*., 1996). However, halothane (Ahmed et al 2020) and pentobarbital (Moghadam and Winlow, 2019) have also been found to mobilize calcium from internal stores in cultured *Lymnaea* neurons: halothane can do this in the absence of external calcium, but pentobarbital cannot, presumably because it modifies the lipid-raft-protein interactions in the cell membrane, (Sierra-Valdez et al, 2016) thereby depressing calcium entry

Depression by anaesthetics of I_Ba_ together with observed changes in activation and inactivation could be related macroscopically to channel opening probabilities or to alterations in single channel kinetics, and this cannot be discerned from whole cell recordings. In ω-conotoxin sensitive and DHP resistant calcium channels expressed from human cortical RNA, both the rate of inactivation of open channels, as well as the proportion of channels inactivated, were increased by barbiturates (Gundersen *et al*., 1988). The changes in both activation and inactivation rates of I_Ba_ argues against a simple blocking mechanism or a decrease in the number of channels available for activation. This study confirms that anaesthetic agents can accelerate calcium current decay, shown previously, for example in pedal I cluster neurons of *Lymnaea* for halothane (Yar and Winlow, 2016); in *Helix* for barbiturates (Nishi & Oyama, 1983); and for halothane in various vertebrate cells, including clonal pituitary cells (Herrington *et al*., 1991), rat sensory neurons (Takenoshita & Steinbach, 1991); hippocampal CA1 neurons (Krnjevic & Puil, 1988) and bovine adrenal chromaffin cells (Charlesworth *et al*., 1994). Depression of calcium current by antagonists belonging to the dihydropyridine (DHP) class of drugs is similarly accompanied by accelerated inactivation (Sanguinetti & Kass, 1984),(Hess *et al*., 1984; Lee & Tsien, 1983) though not invariably (Gross & Macdonald, 1988).

Contrasting with the findings of this study and those of Yar and Winlow (2016), in which volatile anaesthetics produce little or no displacement of the I–V curve along the voltage axis, DHPs tend to shift calcium channel inactivation to more negative potentials. In addition, divalent cation concentrations in the perfusate are known to reduce the potency of these drugs (Lee & Tsien, 1983). Conversely, volatile anaesthetic induced depression of Ca^2+^ influx through L-type and DHP insensitive voltage operated channels is not relieved by increasing [Ca^2+^]_out_ (Gross & Macdonald, 1988), indicating that anaesthetic action is independent of changes in surface potential.

Our results show that a variety of general anaesthetics, including the non-volatile pentobarbitone, produce dose-dependent depression of whole cell HVA Ca^2+^ channel current, I_Ba_. The depression of I_Ba_ is in direct proportion to drug concentration and differences in dose-response between agents is princI_Ba_lly one of magnitude. The efficacy of halothane, isoflurane and sevoflurane to depress I_Ba_ reflects their rank order of clinical potency when expressed as MAC values. Thus, the effect of general anaesthetics on neuronal HVA calcium channels may be fundamental to clinical response.

## Acknowledgments

TJM was partially supported on and EC International Scientific Cooperation Initiative, Contract No. C1*CT93011.

## Notes

### Competing Interest Statement

The authors have declared no competing interest.

